# A bacterial growth law out of steady-state

**DOI:** 10.1101/257709

**Authors:** Yael Korem Kohanim, Dikla Levi, Ghil Jona, Anat Bren, Uri Alon

## Abstract

Bacterial growth depends on numerous reactions, and yet follows surprisingly simple laws that inspired biologists for decades. Growth laws until now primarily dealt with steady-state exponential growth in constant conditions. However, bacteria in nature often face fluctuating environments, with nutritional upshifts and downshifts. We therefore ask whether there are growth laws that apply to changing environments. We derive a law for strong upshifts using an optimal resource-allocation model that was previously calibrated at steady-state growth: the post-shift growth rate equals the geometrical mean of the pre-shift growth rate and the growth rate on saturating carbon. We test this using chemostat and robotic batch culture experiments, as well as previous data from several species, and find good agreement with the model predictions. The increase in growth rate after an upshift indicates that ribosomes have spare capacity. We demonstrate theoretically that spare ribosomal capacity has the cost of slow steady-state growth, but is beneficial in fluctuating environments because it prevents large overshoots in intracellular metabolites after an upshift and allows rapid response to change. We also provide predictions for downshifts for future experimental tests. Spare capacity appears in diverse biological systems, and the present study quantifies the optimal degree of spare capacity, which rises the slower the growth rate, and suggests that it can be precisely regulated.

## Introduction

Systems biology aims to find principles for complex biological phenomena. One way to identify principles is by understanding patterns in biological data. A prime example of such patterns are bacterial growth laws that relate exponential growth rate to cellular and environmental parameters. Although the growth rate *µ* depends on thousands of molecular reactions, it follows surprisingly simple rules. For example, there is a linear relation between *µ* and the ribosomal content of the cell *R* (Ecker and Schaechter, 1963), a law that has been extensively replicated (Scott et al., 2010; Zaslaver et al., 2009), and which inspired foundational mathematical modelling of bacterial resource allocation (Bremer and Dennis, 2008; Churchward et al., 1982; Ehrenberg and Kurland, 1984; Kremling et al., 2007).

The linear relation between growth rate *µ* and ribosomal content *R* also inspired contemporary work by Hwa and co-workers that identified new bacterial laws that connect *µ* and cellular content (Hui et al., 2015; Scott et al., 2010; You et al., 2013). For example, the proteome fraction for carbon utilization, *C*, is a decreasing linear function of growth rate, and *R + C* is approximately constant across growth rates on limiting carbon. These laws generated vigorous research (Bosdriesz et al., 2015; Giordano et al., 2016; Kafri et al., 2016; Maitra and Dill, 2015; Pavlov and Ehrenberg, 2013; Towbin et al., 2017; Weiße et al., 2015), with mathematical models that explain phenomena such as dependence of cellular content on antibiotics (Scott et al., 2014) and the switch between carbon utilization strategies (Basan et al., 2015; Mori et al., 2017a).

Most growth laws until now apply to *steady-state* exponential growth, which occurs when bacteria have been growing for at least several generations in constant conditions (Maaløee and Kjeldgaard, 1966; Shachrai et al., 2010; Wang et al., 2010). In nature, however, bacteria often face changing environments. In particular, they often go from poor conditions with slow growth to richer conditions with more rapid growth, changes known as nutritional upshifts (Poulsen et al., 1995). Despite the considerable experimental and theoretical research going back to Maal0e and Koch on nutritional upshifts (Brunschede et al., 1977; Dennis, 1974; Ehrenberg et al., 2013; Erickson et al., 2017; Giordano et al., 2016; Koch and Deppe, 1971; Maaløee and Kjeldgaard, 1966; Mori et al., 2017b; Pavlov and Ehrenberg, 2013; Sloan and Urban, 1976), no simple upshift growth law was yet described.

Here, we combine theory and experiment to quantify nutritional upshift dynamics. We apply a resource allocation model, which was previously calibrated on steady-state growth experiments (Towbin et al., 2017), and use it to study nutritional upshifts. The model predicts that the growth rate after a large upshift, *µ*_*1*_, is equal to the geometrical mean of the pre-shift growth rate *µ*_0_ and the growth rate on saturating carbon 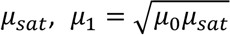. Intuitively, this square-root law stems from an increase in both ribosomal sector and ribosomal saturation level with growth rate. We test the model predictions using chemostat and batch-culture experiments with different carbon sources and temperatures, as well as re-analysis of published data, and find that the square-root law fits the data across conditions, experimental systems and species. This finding precisely quantifies the degree of sub-saturation or spare capacity for growth, showing that ribosomes are sub-saturated in all but the highest growth rates. We propose that ribosomal sub-saturation supplies benefit by i) preventing large overshoots in intracellular metabolites after an upshift and ii) allowing faster growth immediately after an upshift, at the cost of a reduction in steady-state growth rate. Ribosomal sub-saturation, and the upshift growth law, are therefore selectable in environments where upshifts occur frequently.

## Results

### Optimal resource allocation model predicts that the immediate growth rate after large upshifts is the geometric mean of the pre-shift and saturating growth rates

To study upshifts we employ a minimal resource allocation model, which was developed and calibrated for steady-state growth in diverse carbon sources by Towbin et al (Towbin et al., 2017). Here we generalize the model and take it out of steady-state to derive a prediction for growth rate after an upshift.

In the model (Fig. 1a-b), carbon uptake and biomass synthesis are described as a two-enzyme system, composed of a catabolic sector (denoted as the *C* sector) and a ribosomal sector (denoted as the *R* sector). The catabolic sector, which includes carbon transporters and catabolic enzymes, is responsible for carbon uptake and conversion into intracellular substrates (denoted by *x*). The ribosomal sector, which includes ribosomes and translational machinery, take these substrates and converts them into biomass. A third sector (the Q sector, Text S1) includes all other proteins, which under limiting carbon conditions do not change with growth rate (Scott et al., 2014). The experimentally observed tradeoff between making *R* and *C* sector proteins is summarized by *R + C = 1* (You et al., 2013).

**Figure 1:**
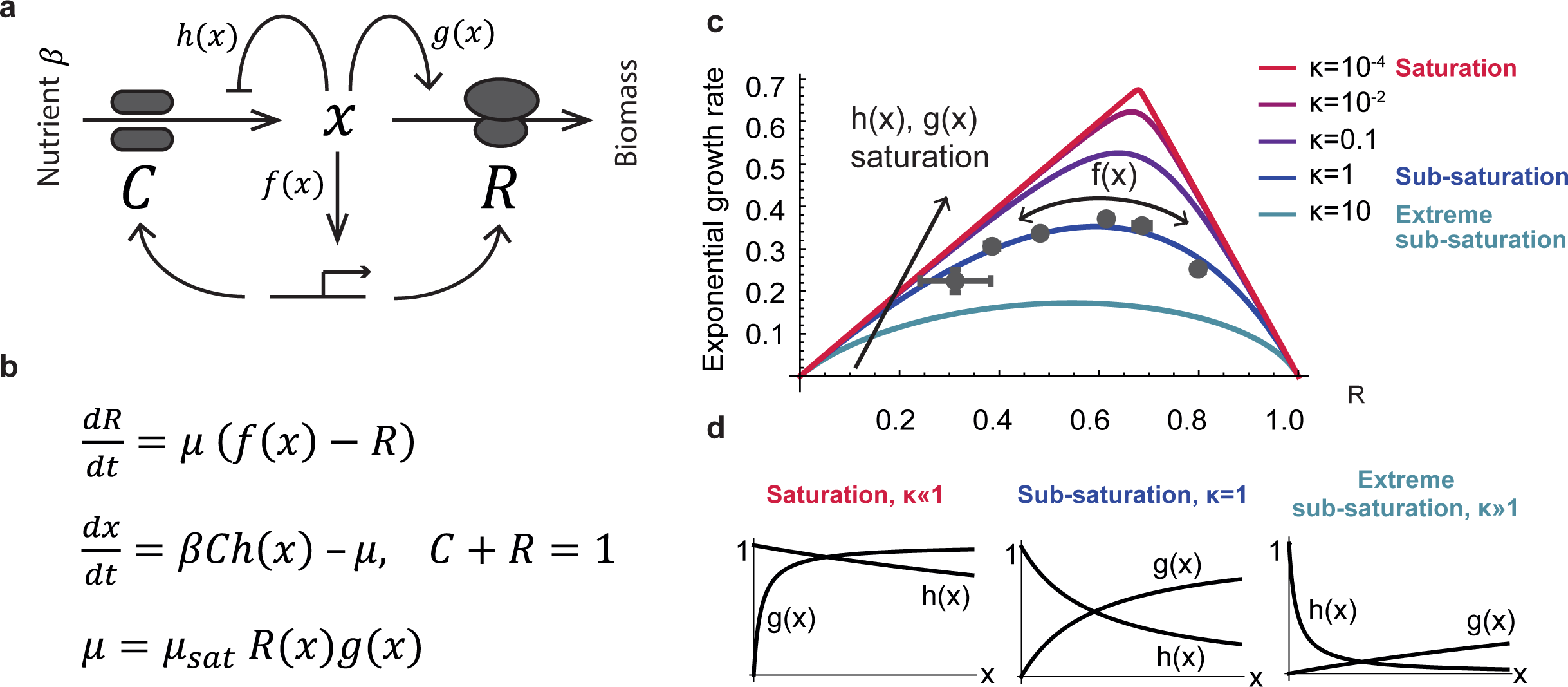
Optimal resource allocation model for bacterial growth. **(a)** Schematic description and **(b)** equations for the cellular resource allocation model. **(c)** Growth curves describe the relation between the allocated ribosomal fraction and the steady-state growth rate in a given environment. Too little or too many ribosomes result in a sub-optimal steady-state growth rate. The optimal regulation function *f(x)* brings steady-state ribosomal content *R* to its optimal value in terms of growth - the maxima of the curves. Different *h(x),g(x)* offset the curve, with a tent-like curve with high optimal steady-state for functions close to saturation (red curve) and a shallow rounded curve with low optimal steady-state growth rate for sub-saturated functions (blue curve). We use MM functions *h(x) = k*_1_*/(k*_1_ *+ x), g(x) = x/(k*_2_ *+ x)*, and measure saturation level by κ*= k*_2_*/k*_1_ (Text S4). Small ***κ*** values reflect a regime where both transporters and ribosomes work close to full saturation for a large range of substrate levels (red pointy curve in **c**, left bottom panel in **d**). *κ* = 1 corresponds to sub-saturation of transporters and ribosomes (purple curve in **c**, middle panel in **d**). *κ* ≫1 leads to extreme sub-saturation with a large substrate range in which both ribosomes and pumps are not efficient (blue shallow curve in **c**, right panel in **d**). We used *β=* 2 to compute all growth curves. Data points are taken from (Towbin et al., 2017), and represent perturbation experiments in which the allocation to catabolic and ribosomal sectors was tuned by externally supplying cAMP to a mutant strain which cannot endogenously produce it (Methods). The data is described best by κ values on the order of 1, suggesting that ribosomes and transporters work at sub-saturation. These experiments suggest that while optimality is reached within a given curve, the curve itself is not optimized for steady-state growth. Error bars are STE of 3 day-day repeats (sometimes smaller than marker).

The exponential growth rate *µ* is the product of the ribosomal sector size *R* and the average rate of the ribosomes, ***g.*** Both *R* and *g* depend on intracellular substrates *x:*

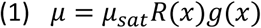

*g(x)* and *R(x)* are normalized between zero and one, such that *µ*_*sat*_ is the growth rate when *x* is saturated. *g(x)*, the average elongation rate, is an increasing function of *x*, describing ribosome utilization (Dai et al., 2016). A sub-saturated ribosomal sector *g(x) < 1* can be the result of either a differential elongation rate or a fraction of inactive ribosomes - both are equivalent in terms of this model.

The internal substrate *x* dynamics are in turn a balance between utilization for biomass production at rate *µ*, and the import of nutrients by the *C* sector:

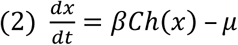

Nutrients are imported and catabolized by the *C* sector at a rate *βh(x)*, where *β* represents nutrient availability. *h(x)*,the import rate, is a decreasing function of *x* which describes inhibition of the transporters by intracellular substrates (Doucette et al., 2011).

The dynamics of *R* are a balance of production and dilution by cell growth:

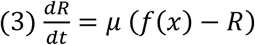

The function *f(x)* describes transcriptional control that determines the fraction of the total biomass production rate *µ* that goes to the *R* sector (Dalebroux and Swanson, 2012). At steady-state, *R = f(x).* For a detailed derivation of the model see SI (Text S1).

A key feature of this model is that growth rate in a given nutrient *β* depends on the control function *f(x)*, because too few or too many ribosomes result in slow growth (Fig. 1c, Text S2). In rich environments (large *β*) more ribosomes are needed, whereas in poor environments more transporters are needed to provide the fastest growth. While the activity curves of the *C* and *R*sector rates, *h(x)* and *g(x)*, define the possible set of steady-states (i.e. the curve in Fig. 1c), *f(x)* determines the chosen steady-state among this set. Importantly, Towbin et al experimentally found that the wild-type level of ribosomes, R, maximizes the steady-state exponential growth rate under many conditions (Towbin et al., 2017). We hence asked whether there exists an optimal *f(x)* that provides the fastest growth for any environment *β*. Using a calculus-of-variations approach (Text S3), we found that indeed such an optimal *f(x)* exists:

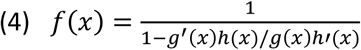

Intuitively, this optimal regulation function *f(x)* determines the best trade-off between *R* and *C*, by balancing the relative advantage of investing in each of these sectors according to the logarithmic sensitivities of their activity curves *h’/h* and *g’/g* (Rosenheim et al., 2010).

With the optimal *f(x)* of Eq. (4), cells are guaranteed to find the optimal growth rate for any nutrient *β.* However, the value of the optimal growth rate can change for different choices of *h(x), g(x)*, which set the transporter and ribosome activity at a given substrate level. In particular, when ribosomes and transporters are fully saturated (*g = h =*1), growth rate is higher than if they are unsaturated (compare tent-like curve to the more rounded curves in Fig. 1c).

Towbin et al calibrated the model for steady-state growth in different carbon sources, using Michaelis-Menten-like (MM) saturation curves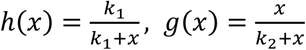. The halfway saturation points for pumps and ribosomes are *k*_1_and *k*_2_respectively. The ratio between these halfway coefficients κ=k_2_/k_1_ represents the cellular saturation level (Fig. 1d, Text S4). Full saturation, in which ribosomes and transporters work close to saturation in a wide range substrate levels, is captured by κ≪1 (Fig. 1d left). Extreme sub-saturation in which ribosomes and transporters work far from their full capacity in most conditions means κ≫1(Fig. 1d right). Medium sub-saturation in which the halfway coefficients of ribosomes and transporters are equal is captured by κ=1(Fig. 1d middle). Towbin et al experimentally manipulated the C sector and measured the resulting growth rate as in Fig. 1c. These experiments indicated that the saturation halfway points are approximately equal, *k*_1_ = *k*_2_, and hence κ~1.

Here, we take the model out of steady-state, and use it to study upshifts. Upshifts are modelled by increasing nutrient availability *β* from a low to a high value. We use the parameter κ=1. We also model full saturation and sub-saturation by varying κ.

The model allows us to calculate the growth rate before the upshift, *µ*_0_, and immediately after the shift *µ_**1**_* (Fig. 2a). This yields a relation between the normalized post-shift and pre-shift growth rates 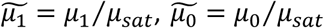 for large upshifts (upshifts into saturating carbon, such that long after the upshift, the growth rate is *μi*_sat_, Text S4), as a function of **κ** (Fig 2b):

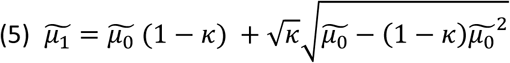

**Figure 2:**
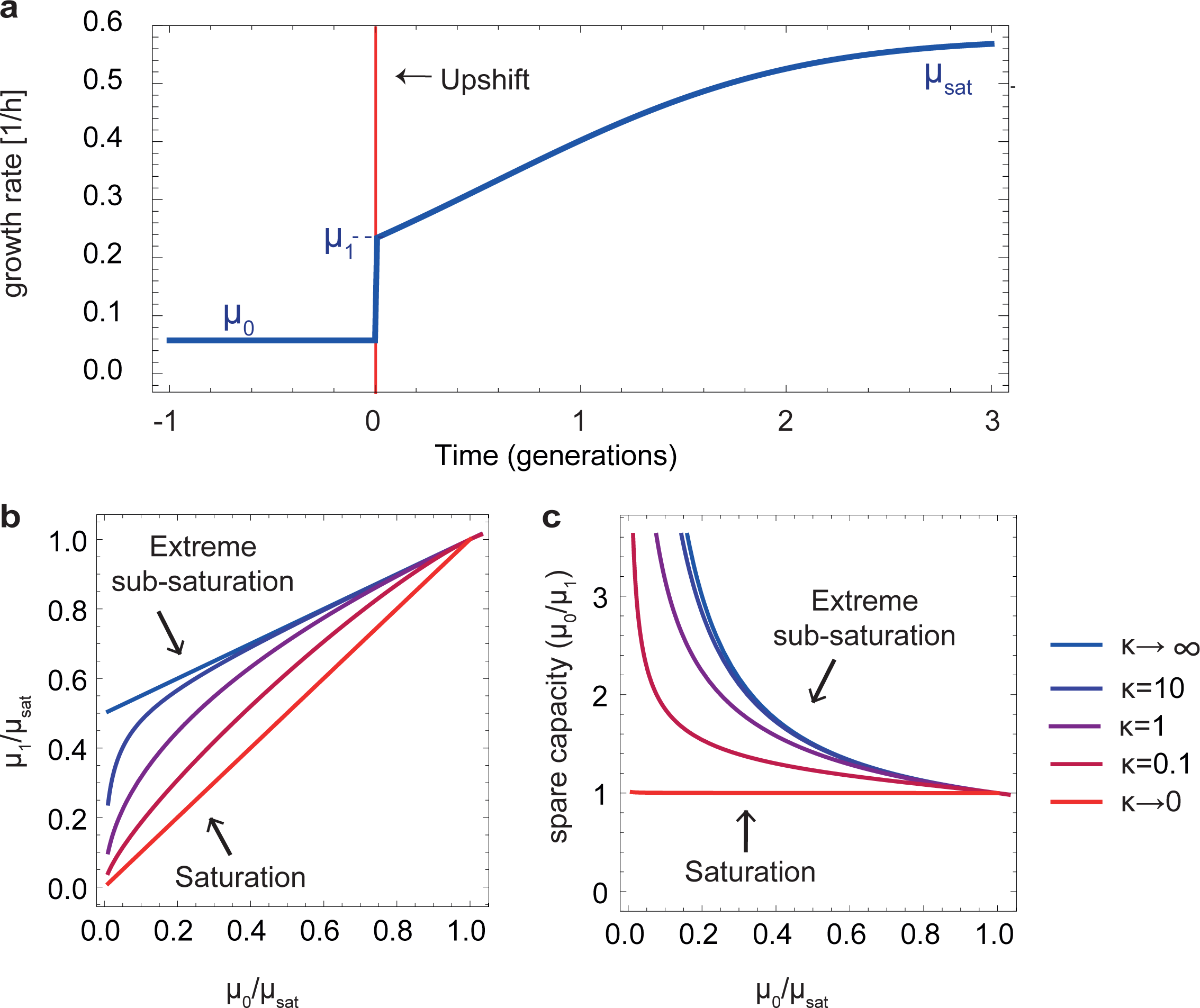
The model provides predictions for growth rate after an upshift. **(a)** Typical growth rate dynamics during nutritional upshift. The growth rate before the shift is *µ*_0_. Upon addition of a rich carbon source (red line), the growth rate rises within minutes to reach a new value *µ*_*1*_, and then slowly increases on the time scale of hours until it reaches its new steady-state value *µ*_*sat*_. The growth rate dynamics were computed from the model with parameter values *η*_0_ = 0.1, *η_**1**_ = 10, k* = 1 (Text S8). **(b),(c):** Model predictions with different k values, which correspond to different levels of saturation for the normalized growth rate. **(d),(e):** Saturation level of ribosomes and transporters as function of the growth rate for different k values.

At full saturation, *k*≪1, the expression reduces to 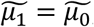, because when ribosomes are saturated before the shift, growth rate cannot immediately increase after the upshift. At extreme sub-saturation, κ≫1, the model gives a linear relation between the normalized preshift and post-shift growth rates with an intercept of ½ (Fig. 2b, dashed lines).

When κ=1, as suggested by the findings of Towbin et al., this expression yields a simple prediction: the Immediate growth rate after an upshift is the geometric mean of the pre-shift and saturating growth rates,

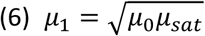

Intuitively, this square-root law results from the following situation: in poor conditions both ribosomal content R and saturation level g are a small value ϵ, and the growth rate is *µ*_0_~ϵ^2^,whereas soon after the upshift, ribosomes are still *ϵ* but saturation is high due to the presence of nutrient, resulting in *µ*_*1*_*~ϵ.* In supplementary note S5 we relax the assumption that h(x) and g(x) are MM-like, and derive a similar square-root law for general h(x), g(x) functions (Text S5).

In the case *κ =*1, the upshift law can be expanded to include general upshifts and downshifts (Text S6), not only large ones as assumed above, resulting in the formula:

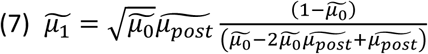

Where the growth rate far after the shift is denoted 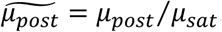. In the case of large upshifts to saturating carbon, 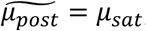, and the formula reduces to the square root formula of Eq. 6 (Text S6).

The upshift law can be recast in terms of the spare capacity of the cells for growth (Diamond, 2002). Cells grow at and then jump to *µ*_1_, indicating that they were operating below full capacity. The spare capacity U can be defined as the fold-change in growth rate after the shift, 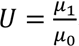. This definition, together with Eq. (6), leads to a spare capacity of

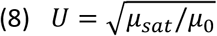

as shown in (Fig. 2c). Spare capacity is smallest (U=1) when cells are close to their saturating growth rate. Spare capacity increases the poorer the medium (the smaller *µ_0_/µ_sat_*). For example, in a medium that allows only 10% of the growth rate on saturating carbon, *µ*_*sat*_=0.1, the cell grows 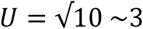 times faster when shifted to saturating carbon.

The present model predicts that at zero growth rate *µ*_0_= 0 there is no upshift, *µ*_1_= 0. This cannot capture the irreducible ribosomal fraction of the bacterial proteome that allows recovery from stationary phase (Dai et al., 2016; Madar et al., 2013). The model can be extended to include such an irreducible ribosomal fraction (Text S7).

### Experimental tests for nutritional upshifts support the square root formula

To test the model predictions, we carried out experiments in which *E. coli* MG1655 cells were shifted from exponential growth in a poor medium to saturating glucose medium. We used two experimental systems: (i) a chemostat, in which slowly dividing cultures in glucose-limited medium (0.02%) were shifted to growth in 0.2% glucose (Fig. 3a, Fig. S1). (ii) Batch culture in a multi-well robotic assay at several temperatures (25°C, 30°C, 37°C) in which cultures growing exponentially on different carbon sources (acetate, sorbitol, rhamnose or pyruvate) were shifted to 0.4% glucose medium (Fig. 3b, Fig. S2,S3). Because we are interested in biomass growth rate, we measured the optical density (OD) of the cells at a temporal resolution of 0.5 min in the chemostat and 3.6 min in the batch culture, with an error of 4-10% in growth rate between biological repeats.

**Figure 3:**
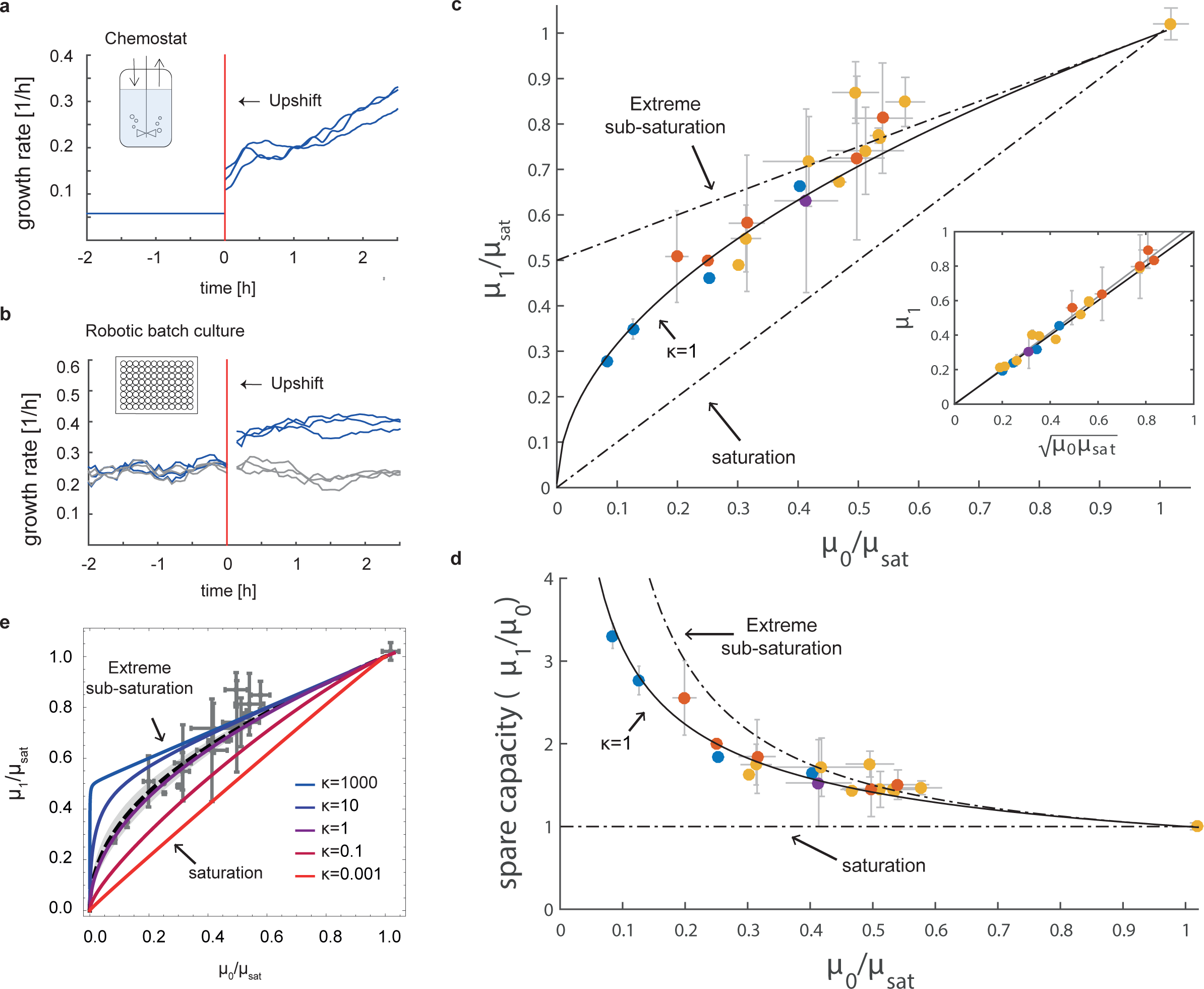
Experimental data support the upshift growth law. **(a)** Growth rate in a chemostat upshift experiment. Cells in limiting-glucose M9 minimal medium (0.02%) with varying doubling times (here 4h) were shifted into high glucose medium (0.2%) (red line). The curves represent 3 biological repeats. **(b)** Growth rate in a multi-well batch culture upshift experiment. Exponentially growing cells on M9 minimal medium with a poor carbon source (here acetate) were either supplemented with high glucose medium (0.4%) (blue curves) or with the pre-shift medium as a control (gray curves). The red line marks addition time. Curves are 3 biological repeats. Growth rate was computed from a window of time-points (Methods), precluding an accurate estimate for about 9 min after the shift, resulting in a gap in the plot. **(c)** Post shift growth rate as a function of pre-shift growth rate is well described by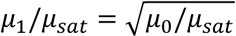(black line), which corresponds to Eq. 6. Dashed lines - model predictions when ribosomes and carbon transporters work at full saturation (*k* ≪ 1) or very far from saturation (*k*≫2). Yellow dots - robotic batch culture experiments, blue dots - chemostat experiments (error bars are STE of 3 day-day repeats repeats), red dots - data from (Maaløe and Kjeldgaard, 1966)/(Sloan and Urban, 1976). Purple dot - yeast data from (Metzl-Raz et al., 2017) (error bars described in SI). For a detailed description of the data points see Fig. S6, Table S1. Inset: data plotted versus Eq. 6. (d) Spare capacity for growth computed form the data, and from the model with *k* = 1(full line), and with high and low saturation (dashed lines). **(e)** Comparison of data (gray points) with model for different values of k. Model with the best fit *k* = 1.3 is in heavy dashed line, with 95% confidence intervals in gray.

We find that growth rate increased abruptly after the upshift by up to 3.3-fold, starting from an initial value *µ*_0_ before the shift and stabilizing after about 15-30 minutes at a new value *µ*_1_ (Fig. 3a,b, Fig. S1, Fig. S3). No such increase was found in control experiments in which pre-growth medium was added instead of glucose, or in the case where cells were shifted from saturating glucose to higher levels of glucose. After the initial increase, the growth rate further increased more slowly on the timescale of hours. We also measured the growth rate on saturating carbon, *µ*_sat_, defined for a given medium and temperature as the exponential growth rate in batch culture with saturating glucose (Fig. S4).

The rapid increase in growth rate from *µ*_0_ to *µ*_*1*_ cannot be explained by synthesis of new ribosomes (Koch and Deppe, 1971). As suggested by Koch and others (Harvey, 1973; Koch, 1988), this hints that cells in slow growth have a higher translational capacity than is actually being used.

In addition to the experiment performed here, we collected data from previous studies on a different strain, *E. coli* 15T^−^, on a different bacterial species, *S. typhimurium*(Maaløe and Kjeldgaard, 1966; Sloan and Urban, 1976), and on the yeast *Saccaromyces cerevisae* (Metzl-Raz et al., 2017). In these experiments, cells were transferred from various carbon sources (fumarate, succinate, aspartate, glyoxylate, galactose or glycerol) to rich carbon sources (saturating glucose or broth), corresponding to strong upshifts. Growth rate was measured by radioactive amino acid incorporation (Maaløe and Kjeldgaard, 1966), OD measurements (Sloan and Urban, 1976) or microscopy (Metzl-Raz et al., 2017) (Methods). Together, the different data sources span a range of conditions, growth rates (0.07 - 1.67 h^−1^), strains and measurement methods (Table S1).

Analyzing the results for the pre-shift growth rate *µ*_0_ vs. the post-shift growth rate *µ*_*1*_ did not reveal a clear pattern, because the same pre-shift rate *µ*_0_could result in different post-shift rates *µ*_*1*_ depending on the conditions (Fig. S5). However, a data collapse occurred when taking into account the condition-dependent value of the saturating growth rate, *µ*_*sat*_ (Fig. 3c) As predicted by the model, the data is well-described by Eq. (6): the growth rate right after an upshift is well described by the geometric average of the pre-shift and saturating growth rates (Fig. 3c,d, Fig. S6, Table S1, Pearson correlation=0.99, p-value <10^−26^).

We also compared the data to the model with different values of *κ.* Fitting equation (5) to the data, we find that the best-fit saturation level *κ* is 1.3 (0.7, 2.5 95% confidence interval, Fig. 3e, Text S4), providing independent support to the steady-state evidence of Towbin et al that the halfway coefficients of ribosomes and transporters are similar.

We also tested alternative mathematical relationships for the data, such as linear regression 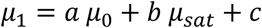 as described in the SI. Such relationships have free parameters, whereas Eq. (6) is a fit with no free parameters. Despite these free parameters, the best-fit error of the alternative formula is comparable or higher than Eq. (6) (R^2^, Table S2). Future experiments with lower error can further test the present predictions.

### Ribosome spare capacity prevents large substrate overshoot, and is beneficial in frequently changing environments

The experiments indicate that ribosomes at low growth rates work far from saturation, because growth increases abruptly after the upshift (Erickson et al., 2017; Mori et al., 2017b). This subsaturation raises a question, because steady-state exponential growth rate is maximal when ribosomes are saturated (Fig. 1c). Maximum growth rate is the reason why many previous models assumed ribosomal saturation (Neidhardt, 1999).

We hypothesize that there are evolutionary benefits to ribosome sub-saturation. The first potential benefit is that sub-saturation of ribosomes and pumps prevents large overshoots of the carbon intermediate x upon upshifts. Such overshoots can be toxic due to osmotic and hydration effects. The saturated model shows an overshoot of tens to thousands of folds in x after an upshift, because the ribosomes have no spare capacity to process the excess carbon, and the pumps cannot reduce their rate effectively to reduce influx. Since many metabolites in central carbon metabolism are in the mM range (Albe et al., 1990), an overshoot of 1000 would raise them to the 1M range which is biologically unfeasible. In contrast, the unsaturated model shows only a small (e.g. ten-fold) transient increase in x upon upshift (Fig 4a,b).

**Figure 4:**
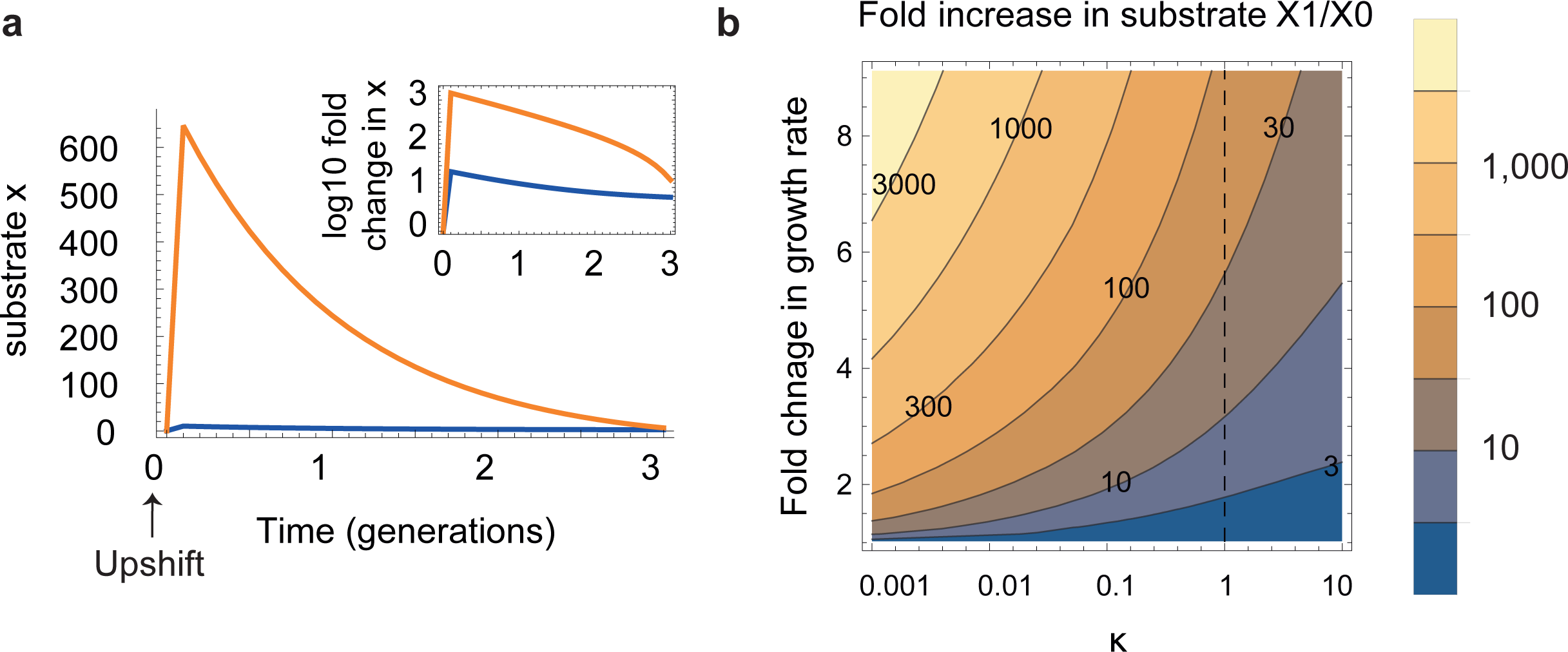
Model predicts that sub saturation of ribosomes avoids large overshoots in metabolic intermediates after an upshift. **(a)** Model results for the substrate x after an upshift (increase in *β* from 0.2 to 5, corresponding to a change in steady-state growth rate of 4 fold, similar to a shift from acetate to glucose). A large overshoot is seen for saturated ribosomes (*k* = 10^−3^), and a much smaller overshot for *k* = 1. Inset: log of the change in x (log10(x(t)/x(0)) where x(0) is the pre-shift steady state value. **(b)** Substrate overshoot after an upshift, relative to pre-shift steady state, max(x(t))/x(0), as a function of ***k*** and the upshift strength (relative change in steady state growth rate).

A second benefit of spare capacity is a growth advantage at early times after an upshift (Mori et al., 2017b). Saturated ribosomes do not allow an increase in growth rate right after an upshift, and result in *µ*_*1*_ *= µ*_0_. This is because all ribosomes are already working full speed before the shift, and an increase in growth rate cannot immediately occur but rather requires synthesis of new ribosomes (Koch, 1988). Sub-saturation therefore has an advantage when upshifts occur often: the benefit of increased growth rate after the shift can offset the cost of lower exponential growth rate at steady-state. In contrast, in conditions in which upshifts are rare, cells with saturated ribosomes have an advantage due to their higher steady-state growth.

To test this hypothesis, we simulated batch growth in which cells with saturated and unsaturated ribosomal strategies competed over limiting nutrient (Fig. 5, Text S8). The model allows us to simulate bacteria with different amounts of spare capacity (sub-saturation), by introducing different values of 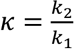.

**Figure 5:**
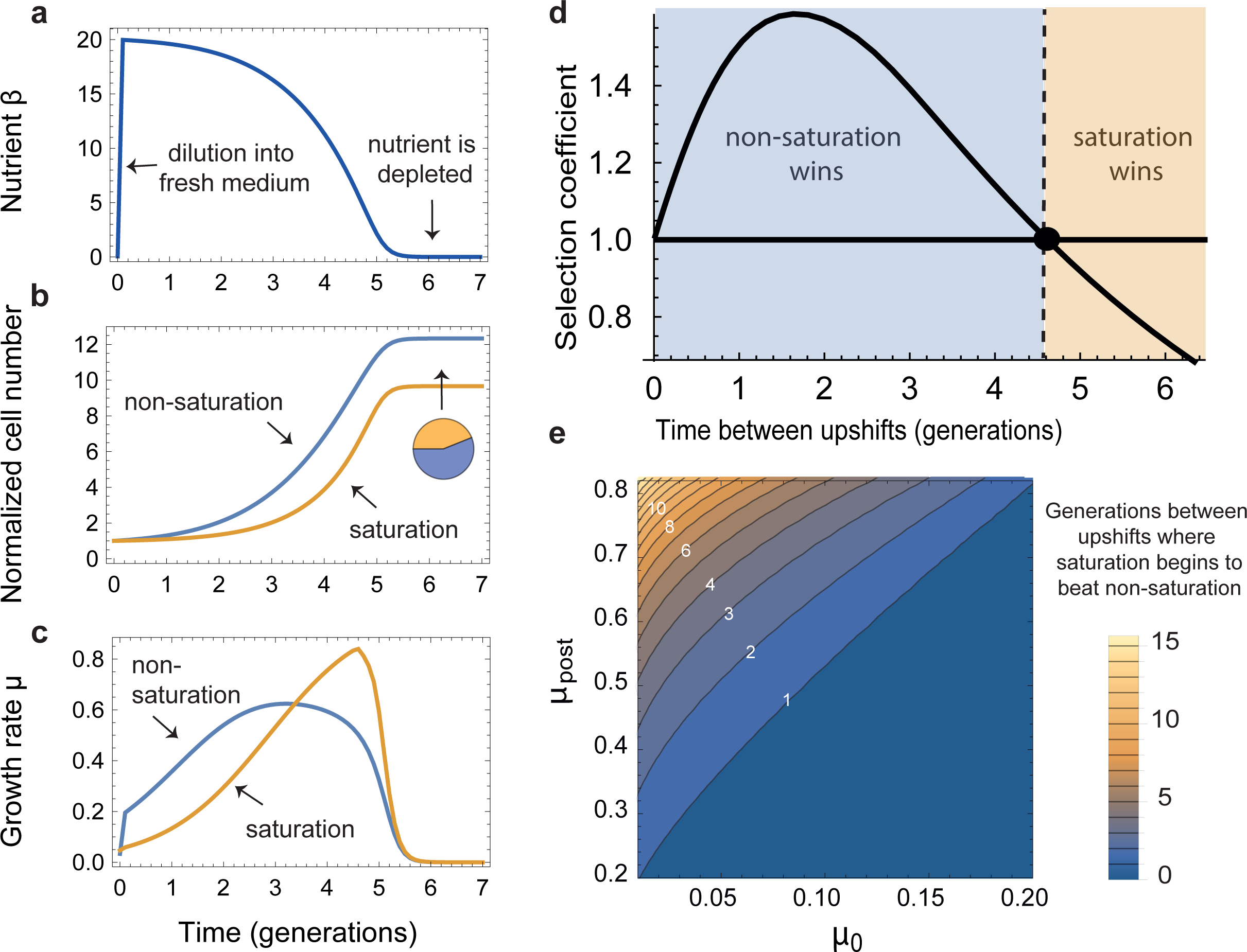
Competition simulations show that sub-saturation is selectable in sufficiently fluctuating environments. **(a)** In nutritional upshift simulations, two populations, one with subsaturated ribosomes (*k* = 1. blue curve in middle and bottom panels), and one with saturated ribosomes (*κ* = 10^−3^, orange curve in middle and bottom panels), were co-diluted into fresh medium. The two populations competed over the shared limited nutrient resource and depleted it at a rate proportional to their number times their growth rate (top panel). The proportion between the two populations was estimated after stabilizing at a constant value (pie diagram, middle panel). **(b)** This was repeated for different initial cell concentrations, which determined the number of generations of growth until nutrient runs out (typical time between upshifts). The threshold number of generations between upshifts for which saturation and sub-saturation are equally beneficial is marked with a black dot. **(c)** This threshold was computed for different initial and psot-shift steady-state growth-rates.

In the simulations, the two populations *(κ* =1 and *κ*=0.001) were diluted into fresh medium resulting in an increase in nutrient availability β (Fig. 5a), and competed over the nutrient. As cells grew, they depleted the nutrient at rate proportional to their number times their growth rate (Jacques Monod, 1949). As cells consumed the nutrient, growth rate declined to zero, simulating limiting nutrient conditions (Bren et al., 2013). The proportion between the two populations was estimated after stabilizing at a constant value (Fig. 5b). We repeated this for different initial cell concentrations (different initial dilutions), which determined the number of generations of growth until nutrient runs out.

As expected, we found that at steady-state, higher growth rate is always achieved by using the saturation strategy. However, after the dilution, non-saturation achieves higher growth for the first few generations (Fig. 5c). After these few generations, the saturation population managed to catch up and achieved faster growth rate again. The relative advantage of the early gain and the long-term loss in growth depends on the number of generations of growth that the cells went through. For example, for a typical parameter set, non-saturation wins over the saturation strategy (selection coefficient>1) when the number of generations until nutrient runs out is smaller than 6 (Fig. 4d, for *µ*_0_= 0.06*µ_sat_,µ_st_ =* 0.7 *µ*_*sat*_, where *µ* is computed with ***κ =*** 1. See Fig. 4e for other parameters). Due to the exponential nature of the growth, even the transient advantage given by the spare capacity has long-lasting implications 6 generations later.

To generalize these findings, we performed a parameter scan to find the optimal level of spare capacity as a function of the frequency of upshifts in the environment, and on the strength of the upshift (difference between pre and post steady-state growth rates). We find that using full saturation *(κ* ≪ 1, zero spare capacity) is beneficial in environments in which upshifts are rare and mild (small difference between pre and post media). In contrast, the more frequent the upshift and the stronger it is, sub-saturation is more beneficial.

An experimental finding by (Gyorfy et al., 2015) supports this prediction. Gyofry et al compared strains deleted for some of the ribosomal RNA operons (Δrrn) to wild type strains in chemostat and batch culture. The Δrrn strains outcompeted wild-type strains in a chemostat but not in batch culture conditions. We Interpret these findings in the light of the present model: the Δrrn strains have fewer ribosomes (Gyorfy et al., 2015) and hence ribosomes are more saturated. They do better under steady-state conditions (chemostat) due to their higher steady-state growth rate. But Δrrn stains do worse after an upshift (shift from overnight to fresh batch culture), due to the predicted benefits of sub-saturation in the wild-type strain.

A further implication of the model is a growth law for nutritional *downshifts.* Like upshifts, downshifts are prevalent in nature, since bacteria tend to exhaust their nutrient resources during the last generations of exponential growth (Bren et al., 2013). In very rich environments, ribosomes work near saturation, but the transporters are sub-saturated. In downshifts, the model suggests that it is the sub-saturation of *transporters/utilization systems* that is advantageous: In rich environments few transporters (small ***C*** sector) are needed, but after the transition to a poor medium, cells need to produce new transporters and catabolic systems (increased ***C*** sector) to reach optimal growth. Moreover they need to produce these transporters despite the low nutrient influx in the poor medium. If transporters were saturated, growth rate would plummet in the poor condition; however, when transporters are unsaturated before the shift, there is spare capacity to prevent a strong decrease (Fig. S7). Thus, transporter sub-saturation is selectable in conditions with frequent downshifts. The model analytically predicts a growth law for downshifts from a rich medium,

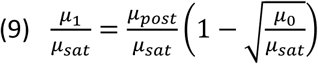

(see Text S6 for the range of validity of this formula, and for a more general formula) where *µ*_*post*_ is the steady-state growth rate in the post-shift medium. Since downshifts are experimentally harder to explore, we defer a test of this prediction to future studies.

## Discussion

We derive a law for growth immediately after an upshift from a model of optimal control of transcriptional regulation in cells. This growth law is supported by experiments in chemostat and robotic batch culture conditions in different media and temperatures, and re-analysis of previous data on *E. coli*, Salmonella and yeast. The growth law is also a quantitative measure of the spare capacity of cells for growth. We suggest that sub-saturation of ribosomes can be beneficial in frequently changing environments, since it prevents large overshoots in metabolic intermediates and allows rapid initial increase in growth rate following an upshift.

The question of whether ribosomes work at saturation has a long history. There seem to be at least two schools of thought. In one school, exemplified by Maaløe and co-workers (Maaløe and Kjeldgaard, 1966), ribosome saturation is almost a law in itself (as for example in a review by Neidhart (Neidhardt, 1999)). This postulate is due to the fact that balanced exponential growth is maximal at ribosomal saturation. The second school of thought suggests that ribosomes are substantially sub-saturated under all but the best conditions, often assuming a fraction of inactive ribosomes. Examples of this way of thinking can be found in work on upshifts by Koch (Koch, 1971, 1988; Koch and Deppe, 1971), Harvey (Harvey, 1973) and subsequent models (Ehrenberg et al., 2013) and experiments (Borkowski et al., 2016; Dai et al., 2016; Metzl-Raz et al., 2017). The present study supports the second school. Importantly, it quantifies the extent of sub-saturation, predicting the spare capacity U for growth as a function of the growth rate *µ*_0_ and the saturating growth rate *µ*_*sat*_, namely 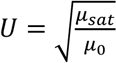 Spare capacity is larger the poorer the medium (the smaller 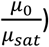

Sub-saturation is selectable in frequently changing environments, because it prevents large overshoots in intracellular metabolites after an upshift which are likely to be toxic. Spare capacity also confers rapid growth after upshifts and downshifts, which may offset the cost of the reduction in growth rate at steady-state. A potential experimental test could compare different bacterial strains and species in terms of their upshift performance and their ribosome saturation. Our model predicts that the higher the saturation, the slower the post-shift improvement, a prediction which is supported by experiments on rrn deletion strains (Gyorfy et al., 2015). This hints at a tradeoff situation (Klappenbach et al., 2000; Shoval et al., 2012; Weiße et al., 2015), in which higher steady-state growth comes at the expense of rapid response to changes. Mori et al., in a paper published during the publication process of this study, reached a similar conclusion (Mori et al., 2017b), elegantly showing how the basal fraction ribosomes is optimal for a given frequency of environmental change.

More generally, metabolomic experiments indicate that sub-saturation is the norm for many metabolic enzymes, which under typical conditions work well below their maximal rate (Davidi et al., 2016). This sub-saturation is often thought of as a “safety factor”, which is beneficial when enzymatic load is uncertain (Diamond, 2002). It would be interesting to check in other systems whether the present quantitative relationship for spare capacity is found (e.g does spare capacity increase at low steady-state system flux or activity (Suarez et al., 1997)) or whether other types of laws govern spare capacity in each context.

The upshift growth law can be tested in additional strains, organisms and conditions. The model predicts specific forms for the ribosomal and transporter saturation and regulation functions which can in principle be tested experimentally. More generally, this study suggests that growth laws can be found for dynamic situations, not only for steady-state growth, deepening our understanding of how bacterial growth dynamics work and what tasks they evolved for.

